# The Role of Paraventricular Nucleus of Thalamus in Sleep Disturbance Induced by Withdrawal from Repeated Ethanol Exposure

**DOI:** 10.1101/2025.06.04.657945

**Authors:** Aubrey Bennett, Jeyeon Lee, Hyunjung Kim, Vatsala Kapoor, David Jury, Seungwoo Kang

**Author notes:** Contributed equally to this work (Aubrey Bennett, Jeyeon Lee). Corresponding author: Seungwoo Kang, PhD, Department of Pharmacology and Toxicology, Medical College of Georgia, Augusta University, Augusta, Georgia 39012, USA.

## Abstract

Sleep disturbance is known to be comorbid with withdrawal from repeated ethanol exposure and could be a negative reinforcement for the majority of people with alcohol use disorder (AUD). The paraventricular nucleus of the thalamus (PVT) has been highlighted for its function in integrating arousal states and associated modulation in sleep homeostasis. However, there is limited understanding of the involvement of PVT neurons in regulating sleep patterns, especially during withdrawal from chronic ethanol exposure. In this study, we investigated the potential function of the PVT in sleep disturbance during ethanol withdrawal using electrophysiology, in vivo calcium imaging, biochemical, and chemogenetic approaches. At 24 hours post-withdrawal from chronic intermittent ethanol exposure (CIE) for four weeks, there is an increase in wake time and a decrease in non-rapid eye movement (NREM) sleep. The calcium transient levels in the PVT neurons are positively correlated with the transition from sleep to wakefulness. CIE elevates the PVT neuronal activity in a subregion-specific manner, resulting in a significant rise in cFos levels in the anterior PVT (aPVT). Temporal suppression of aPVT excitatory neurons via chemogenetics ameliorates the disturbance in sleep patterns generated by CIE. The aPVT has a notable distinction in the expression of the m-type potassium channel subunit, KCNQ2, with a higher expression level compared to the posterior PVT (pPVT). While the expression of KCNQ2 in the aPVT is reduced in CIE mice, the restoration of KCNQ2 expression using viral gene transfer within the aPVT alleviates the sleep disturbances produced by CIE. This data indicates a significant role of the PVT in sleep disturbance during ethanol withdrawal, which may partially be due to the downregulation of M-channels, hence underscoring M-channels in the PVT as a potential therapeutic target for sleep disturbance in alcohol use disorder.

## Introduction

Sleep is essential for maintaining healthy living patterns (Harding and Feldman, 2008). Consequently, disrupted sleep has been linked with various psychiatric disorders, including alcohol use disorder (AUD) (Koob and Colrain, 2020). Clinically, a reciprocal association exists between sleep and alcohol use. While AUD patients frequently encounter sleep difficulties during both initial and prolonged recovery, those with AUD and sleep disturbances face a heightened risk of relapse into alcohol consumption (Koob and Colrain, 2020; Mikl et al., 2025). Furthermore, beyond alcohol-related diseases, sleep disturbances may also contribute to mental and physical health problems. Sleep disturbances increase the risk for mental disorders such as depression, anxiety, and substance misuse, and could be a significant factor in risk of obesity, diabetes, hypertension, and critical vascular disease including stroke and myocardial, which have been spotlighted these days as a risk factor of age-dependent disease like Alzheimer’s, suggesting that sleep disturbances could serve as the initial catalyst for numerous comorbid psychiatric and neurological disorders raised by AUD (Ju et al., 2017; Koob and Colrain, 2020). Thus, it is crucial to have a comprehensive understanding of how sleep is modulated and how sleep patterns are disrupted during alcohol withdrawal to diminish sleep-related social burden. However, the fundamental cellular and molecular mechanisms contributing to sleep disturbance, especially occurring during withdrawal from repeated ethanol exposure, remain unclear.

The paraventricular nucleus of thalamus (PVT), a small dorsal thalamic brain area, has been implicated in controlling various aspects of arousal, including sleep-wake states, attention, and emotional psychological responses (Bu et al., 2022). It receives input from several brain regions including monoaminergic centers such as locus coeruleus, dorsal raphe, and zona incerta for norepinephrine, serotonin, and dopamine, respectively (Sugiyama et al., 2019; Ye et al., 2023), and projects to areas associated with emotional processing, including the limbic system (Zhou et al., 2022). The PVT has recently received attention due to its integrated role in physiological and pathophysiological behaviors involving cognition, reward, emotion, pain, arousal, and sleep (Penzo et al., 2015; Pandey et al., 2019; Barson et al., 2020; Pandey and Barson, 2020; He et al., 2024; Jiang et al., 2024; McDevitt et al., 2024; Koita et al., 2025), as well as its involvement in the pathogenic relationship between sleep and psychiatric disorders (Li et al., 2022a; Zhao et al., 2022). Indeed, pharmacological and chemogenetic modulation of PVT neuronal activity affects sleep patterns (Hua et al., 2018; Matyas et al., 2018; Ren et al., 2018; Gao et al., 2020; Ao et al., 2021). Interestingly, cumulative evidence has shown genetically and functionally distinct cell types in the PVT mainly alongside the anterior-posterior axis (Gao et al., 2020; Gao et al., 2023; Shima et al., 2023). Along the anteroposterior axis, numerous cellular components that regulate neuronal excitability, including neuromodulator receptors and ion channels, exhibit distinct expression in the PVT (Shima et al., 2023). Nonetheless, it is still unclear whether the PVT contributes to the changes in sleep patterns after ethanol withdrawal and if this process entails any molecular adaptations according to the heterogenous characters in the PVT neurons.

In the current study, given the essential role of the PVT in sleep-wake states, we attempted to clarify how PVT neuronal activities are adapted during chronic repeated ethanol exposure and consequently integrate the changes in sleep patterns using a combination of chemogenetic, electrophysiological, pharmacological, genetic, fluorescent-based calcium imaging, and behavioral approaches. Our data suggests that PVT M-channels, one of the potassium channels regulating neuronal firing, could be a potential therapeutic target for sleep disturbance in alcohol use disorder.

## Materials and Methods

### Animals

All experimental procedures were approved by the Augusta University Institutional Animal Care and Use Committee (IACUC) and conducted in accordance with NIH guidelines. The C57BL/6J mouse line (Catalog no. 000664) was acquired from Jackson Laboratory (Bar Harbor, ME). Mice were housed in standard Plexiglas cages. The colony room was maintained at a stable temperature of 24 ± 1°C and humidity of 60 ± 2%, with a 12-hour light/dark cycle (lights on at 07:00 A.M.). Mice were granted *ad libitum* access to food and water. Male mice were used for all the experiments.

### Stereotaxic surgery for AAV injection

Mice were anesthetized with isoflurane (1-1.5% in oxygen gas) and placed on a rotational digital stereotaxic apparatus (Model 69101; RWD Life Science) while positioned on a heating pad to maintain body temperature. The skull was aligned using a dual-tilt equalizer and holes were drilled at the designated stereotaxic coordinates. To perform viral gene transfer, viruses were infused into the anterior PVT (AP −0.5 mm, ML +/-0.0 mm, DV −3.5 mm from bregma) at a rate of 100 nl/min for 3 minutes using a 34-gauge injection needle (Catalog No. NF34BV; World Precision Instruments) with a micro-syringe pump (Model UMP3). The injected needle was held at the injection site for an additional 10 minutes post-injection. The titer of AAVs injected were 3 × 10^12^ – 2 × 10^13^ GC/ml. pAAV-syn-dLight1.3b was a gift from Lin Tian (Addgene viral prep # 135762-AAV9;

RRID:Addgene_135762)(Patriarchi et al., 2018); pAAV-CaMKIIa-hM4D(Gi)-mCherry was a gift from Bryan Roth (Addgene viral prep # 50477-AAV5; RRID:Addgene_50477). pENN.AAV.CamKII 0.4.Cre.SV40 was a gift from James M. Wilson (Addgene viral prep # 105558-AAV5; RRID:Addgene_105558). AAV5-CaMKIIa-RCaMP2 was a gift from Karl Deisseroth (UNC Vector Core). We administered buprenorphine sustained release (1.3 mg/kg, s.c.; CoVetrus) to alleviate post-surgical pain following stereotaxic surgery.

### EEG-EMG recordings for sleep evaluation

For EEG/EMG recordings, mouse models were prepared as described elsewhere (Barger et al., 2019; Ma et al., 2019; Antila et al., 2022). We prepared an electrode socket with a 2 × 4 pin header: two mini-screw channels for EEG recording and two micro-ring channels for EMG recording. Two mini-screws were implanted in the skull above the frontal (AP: −2 mm; ML: - 1 mm) and temporal (AP: 3 mm, ML: - 2.5 mm) cortices for EEG signal sampling, while two micro-rings were placed in the neck muscles for EMG data acquisition. The electrode socket was attached to the skull with dental cement for recording in freely moving mice. The mice were habituated to the EEG recording chamber for three days before recording to reduce environment novelty. Electrophysiological signals were digitized at 1 kHz (RHD system, Intan technologies) and the signal analysis was conducted by a custom-written code in MATLAB. The sleep stages were semi-automatically scored in 2.5-second epochs using an open-source toolbox, AccuSleep (Barger et al., 2019), followed by manual verification through a custom MATLAB-based graphical user interface.

### In vivo Ca^2+^ signal with fiber-photometry

We recorded the cellular Ca^2+^ and dopamine transients simultaneously in real-time in vivo using fiber-photometry, as previously detailed (Kang et al., 2023; Matthews et al., 2023). Briefly, three weeks post-injection of AAV encoding RCaMP2 for calcium and dLight1.3b for dopamine into the PVT, we implanted an optic cannula into the PVT (AP −0.40 mm, ML +0.0 mm, DV −3.3 mm from bregma). The fluorescent signals were captured at 30 frames per second using a fiber photometry system (Plexon Multi-Fiber Photometry System, Plexon, Dallas, Texas), which included a dichroic mirror, and a lens focused on a photomultiplier tube (PMT) to reflect beams from LEDs for three excitation ranges centered at 560 nm, 465 nm, and 410 nm wavelengths. The collected data were analyzed with MATLAB-based photometry modular analysis tool, pMAT (Bruno et al., 2021).

### Brain slice preparation and Ex vivo electrophysiology

Brain slices containing the PVT region were prepared for electrophysiological recordings, as described (Kang et al., 2020). Briefly, mice were deeply anesthetized by isoflurane inhalation, after which the brain was rapidly extracted and immersed in ice-cold sucrose-based artificial cerebrospinal fluid (aCSF) containing the following (in mM): 87 NaCl, 75 sucrose, 2.5 KCl, 11.25 NaH2PO4, 0.5 CaCl2, 7 MgCl2, 25 NaHCO3, 0.3 l-ascorbate, and 25 glucose, and oxygenated with 95% O2/5% CO2. Coronal brain slices (300-350 μm) were cut with a vibrating compresstome (VF-310-0Z, Precisionary Instruments), subsequently placed in a slice holding chamber and incubated for 30 min at 34°C and kept for at least 1 hour at room temperature (24-25°C) in carbonated (95% O2/5% CO2) standard artificial cerebrospinal fluid (aCSF) containing the following (in mM): 126 NaCl, 1.25 NaH2PO4, 1 MgCl2, 2 CaCl2, 2.5 KCl, 25 NaHCO3, and 11 glucose. Electrical signals were captured using an Axon 700B amplifier, a Digidata 1550B A/D converter, and Clampfit 11.0 software (Molecular Devices). Throughout the experiments, the bath was consistently perfused with warm (32 °C) carbonated aCSF at a rate of 2.0-2.5 ml/min. Patch pipettes (6–8 MΩ) were filled with the solution containing (in mM) 130 K-methanesulfonate, 10 KCl, 4 NaCl, 10 HEPES, 0.5 EGTA, 2 MgATP and 0.2 Na2GTP. The pH was adjusted to 7.2 with Tris-base and the osmolality to 310 mOsmol/L with sucrose. Healthy cells were targeted under a high magnification microscope (at 400X, Nikon FN1 Microscope, Melville, NY) by shape (round, ovoid, and non-swelled plasma membrane). M-currents were recorded in the presence of 1 µM tetrodotoxin, at a holding potential of −20 mV, and evoked by 500-ms repolarizing -40 mV (Bordas et al., 2015; Kang et al., 2017).

### Chemogenetics and drug treatments

JHU31670 was purchased from Hello Bio (J60; Princeton, NJ), noted for its superior blood-brain barrier penetrance and enhanced affinity, potency, and selectivity of DREADDs (Designer Receptors Exclusively Activated by Designer Drugs) (Bonaventura et al., 2019). For the activation of DREADDs, we administered J60 (0.3 mg/kg) or saline intraperitoneally 10 minutes prior to the experiments in mice. The concentration has been demonstrated to have no off-target effects in rodent behavioral tests (Bonaventura et al., 2019; Zhang et al., 2020; Costa et al., 2021; Desloovere et al., 2022; Van Savage and Avegno, 2023).

### Repeated intermittent ethanol exposure

The intermittent ethanol exposure paradigm has been described (Hong et al., 2019). Mice were subjected to ethanol exposure in a vapor inhalation chamber (Morton et al., 2014) for four weeks. Each daily’s cycle included either 16 hours of ethanol vapor (CIE group, BEC 60-80 mg/dL at the end of session) or equivalent counterpart room air (CON group), followed by 8 hours of abstinence in their home cage away from both vaporized ethanol and air. This was repeated daily for four consecutive days, succeeded by three days of abstinence (Fig. 1a).

**Figure 1.**
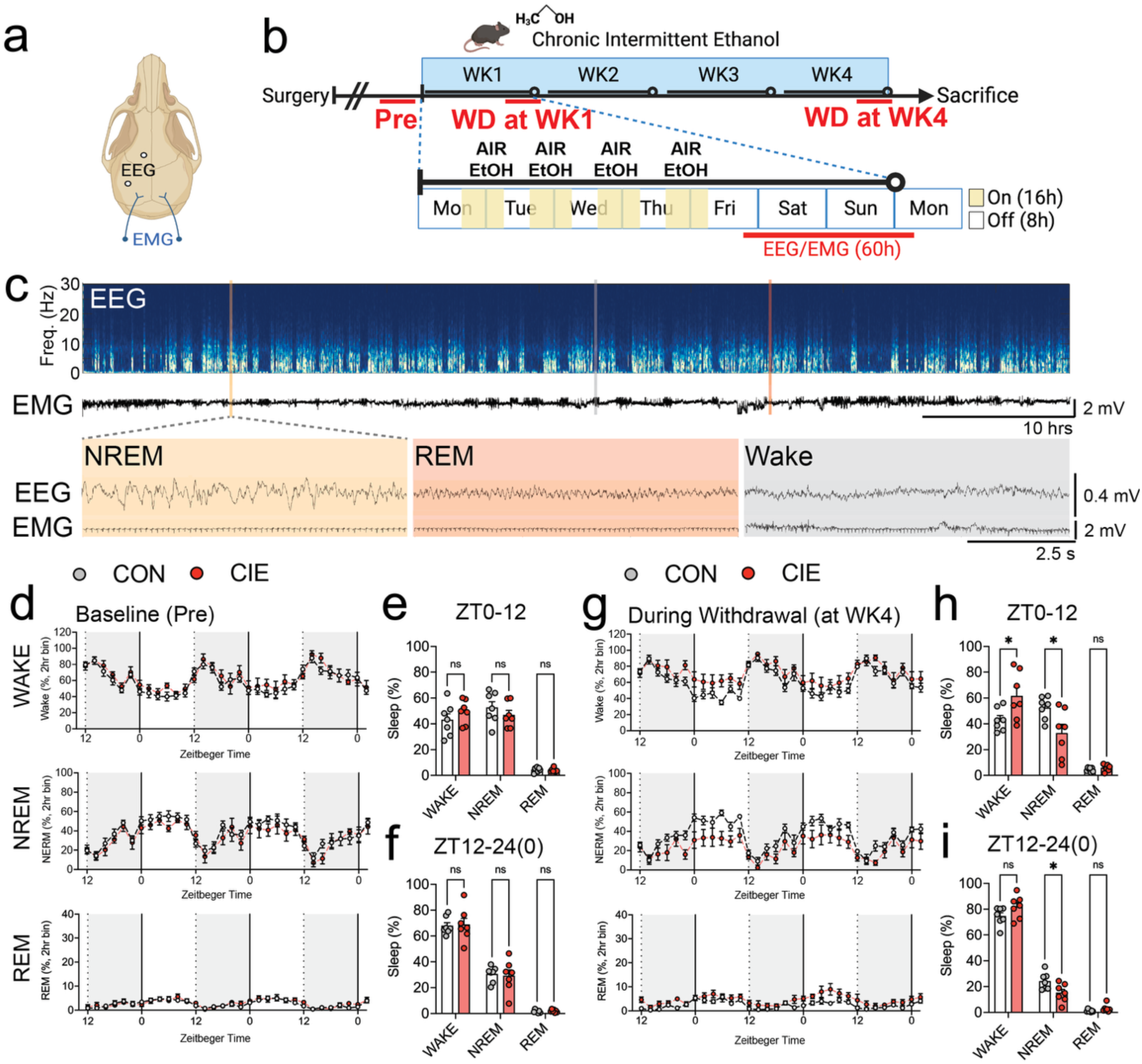
Withdrawal from repeated ethanol exposure induces the disruption in sleep patterns. (a,b) Diagram showing the electrode position and experimental schedule. (c-h) Representative figures (c) and pooled data (d-i) showing that WAKE time is significantly increased during withdrawal from chronic intermittent ethanol exposure for four weeks. [Figure 1e: Two-way ANOVA, Interaction F (2, 36) = 1.558, P=0.2245, Sleep type F (2, 36) = 109.5, P<0.0001, Group F (1, 36) = 3.677e-014, P>0.9999], [Figure 1f: Two-way ANOVA, Interaction F (2, 36) = 0.07375, P=0.9290, Sleep type F (2, 36) = 235.1, P<0.0001, Group F (1, 36) = 4.587e-015, P>0.9999], [Figure 1h: Two-way ANOVA, Interaction F (2, 36) = 8.987, P=0.0007, Sleep type F (2, 36) = 61.47, P<0.0001, Group F (1, 36) = 4.542e-016, P>0.9999], [Figure 1i: Two-way ANOVA, Interaction F (2, 36) = 6.118, P=0.0052, Sleep type F (2, 36) = 579.1, P<0.0001, F (1, 36) = 2.142e-014, P>0.9999]. N=7/group. Two-way ANOVA and Bonferroni’s post-hoc. Data represented as mean ± SEM. *p<0.05.

### Western blotting

Tissue containing the PVT was punched out from coronal slices (1 mm thick) and homogenized in ice-cold RIPA lysis buffer (Thermo Fisher) containing a protease inhibitor cocktail (Roche). Equal amounts of protein extracts were denatured, subjected to SDS-PAGE using 4-12% Bis-Tris gels, and transferred to PVDF membranes (Thermo Fisher). PVDF membranes were blocked with tris-buffered saline containing 0.05% Tween 20 and 5% (w/v) non-fat dried milk. These membranes were incubated with anti-KCNQ2 antibody (1:300, Rabbit, Alomone), and anti-GAPDH antibody (1:2000, Mouse, Sigma Aldrich) overnight at 4 °C. After washing with TBST, the membranes were incubated with appropriate horseradish peroxidase-conjugated secondary antibodies for 1 hour at room temperature. The proteins were visualized by ECL solution (Thermo Fisher) using the G:Box Chemiluminescent Imaging System (Syngene).

### Immunofluorescence

Brains were fixed with 4% paraformaldehyde solution and transferred to 30% sucrose (Sigma-Aldrich) in phosphate-buffered saline at 4°C for 72 hours. Brains were then frozen and sectioned at 50 μm using a microtome (Precisionary). Brain slices were stored at −20°C in a cryoprotectant solution containing 30% sucrose and 30% ethylene glycol in phosphate-buffered saline. Thos floating unfrozen sections were incubated in 0.3% Triton X-100 and 3% bovine serum albumin in phosphate-buffered saline for 1 hour, followed by incubation with primary antibodies in 3% bovine serum albumin overnight at 4°C. Primary antibodies against NeuN (Mouse, 1:500, Abcam), KCNQ2 (Rabbit, 1:300, Alomone) with Goat anti-Mouse IgG AlexaFluore594 (1:500, Abcam) and Goat anti-Rabbit IgG AlexFluore488 (1:500, Abcam) were used. Images were acquired with an LSM 700 laser scanning confocal microscope and Axioscan microscope (Carl Zeiss, Heidelberg, Germany).

### Data analysis

All data are shown as mean ± SEM using Prism 10.4 (GraphPad Software, San Diego, CA). Statistical significance was assessed using paired or unpaired t-test, and One-way or Two-way ANOVA with *post hoc* Bonferroni’s multiple comparisons test, when appropriate. Values of p<0.05 were considered significant.

## Results

### Withdrawal from repeated ethanol exposure induces the disruption in sleep patterns

To determine whether withdrawal from repeated ethanol exposure affects sleep patterns, EEG/EMG was recorded in mice continuously to monitor wakefulness, NREM, and REM sleep repeatedly (Figure 1a,b). After one week of recovery from the EEG/EMG implantation surgery, the mice were exposed to vaporized ethanol or air (CON) via the chronic intermittent ethanol paradigm (CIE) for 16h overnight followed by 8h of abstinence for four days a week. The remaining three days no ethanol exposure was given; this was considered a withdrawal period. This exposure persisted for four weeks with three separate sleep evaluations; the first prior to ethanol exposure as a baseline (Pre), and during withdrawal following week 1 of exposure and week 4 of exposure to better investigate at which withdrawal point sleep disturbance comes into play (Figure 1b). We chose to mainly evaluate the sleep pattens at 24 hours after the ethanol exposure since the timepoint is known to exhibit increased ethanol withdrawal symptoms in rodent models (Perez et al., 2015; Perez and De Biasi, 2015). During this period, we examined non-rapid eye movement (NREM), otherwise known as slow wave sleep, rapid eye movement (REM), and wake sleep patterns (Figure 1c).

During the baseline (Pre) recording, prior to ethanol exposure, there were no significant differences between the CIE and CON groups in the wake, NREM, and REM sleep patterns (Figure 1d-f) [ZT0-12: Two-way ANOVA, F (2, 36) = 1.558, P=0.2245, Pre: CON vs CIE, WAKE: p=0.6174, NREM p=0.7103, REM: p>0.999], [ZT12-24(0): Two-way ANOVA, F (2, 36) = 0.07375, P=0.9290, Pre: CON vs CIE, WAKE: p>0.999, NREM: p>0.999, REM: p>0.999]. While sleep patterns did not change significantly in the group of CIE and counterparts (CON) after 1-week repeated exposure (Figure S1a-c) [ZT0-12: Two-way ANOVA, F (2, 36) = 1.844, P=0.1728, 1WK: CON vs CIE, WAKE p=0.7513, NREM p=0.4349, REM p>0.9999], [ZT12-24(0): Two-way ANOVA, F (2, 36) = 1.903, P=0.1639, 1WK: CON vs CIE, WAKE p>0.9999, NREM p=0.3676, REM p>0.9999], we observed significantly increased percent of wakefulness and reduced percent of sleep, including a reduction in the amount of NREM sleep during the light phase at 4 weeks (Figure 1g,h) [ZT0-12: Two-way ANOVA, F (2, 36) = 8.987, P=0.0007, 4WK: CON vs CIE, WAKE p=0.0196, NREM p=0.0113, REM p>0.9999]. during the dark phase at 4 weeks, there was no difference in wakefulness between groups, NREM was significantly reduced in the CIE compared to this in CON (Figure 1i) [ZT12-24(0): Two-way ANOVA, F (2, 36) = 6.118, P=0.0052, 4WK: CON vs CIE, WAKE 0.1095, NREM p=0.0321, REM p>0.9999]. These results indicate that withdrawal from chronic intermittent ethanol exposure does induce disruptions in sleep patterns and exert wakefulness.

### The neuronal calcium transients in the PVT are positively correlated with the transition of sleep pattern to wake

Previous findings have suggested the PVT as a hub brain area to control sleep-patterns and wakefulness (Colavito et al., 2015; Ren et al., 2018; Zhao et al., 2022; Eacret et al., 2023). Thus, to clarify whether the changes in sleep patterns are related to the neuronal activities in the PVT, we measured spatiotemporal populational Ca^2+^ transients of the PVT neurons, particularly anterior part of PVT (aPVT), with RCaMP2, a genetically encoded fluorescence-based Ca^2+^ indicator, synched with the sleep pattern changes which are validated by the combination of EEG and EMG (Figure 2a and b). We also measured dopamine transients simultaneously, with dLight, a genetically encoded fluorescence-based dopamine indicator, in the PVT, levels of which have been shown to be correlated with wakefulness (Dong et al., 2019). While the PVT neuronal calcium transients were increased during the transition from NREM to WAKE (Figure 2c, Paired t-test, t=4.142, df=17, P=0.0007), the signals were decreased during the transition from WAKE to NREM (Figure 2d, Paired t-test, t=3.766, df=18, P=0.0014). Interestingly, the dopamine transients in the PVT were changed similar to the calcium levels (Figure 2e, Paired t-test, t=5.572, df=17, P<0.0001; Figure 2f, Paired t-test, t=2.620, df=18, P=0.0174), and those were positively correlated with the calcium transients of the PVT neurons during the transitions between NREM and WAKE (Figure 2g, Correlation, R^2^=0.7739, DFn, Dfd = 1, 35, P<0.0001). This data suggests that the enhanced neuronal activity in the PVT is positively correlated with the transition of sleep pattern to wakefulness.

**Figure 2.**
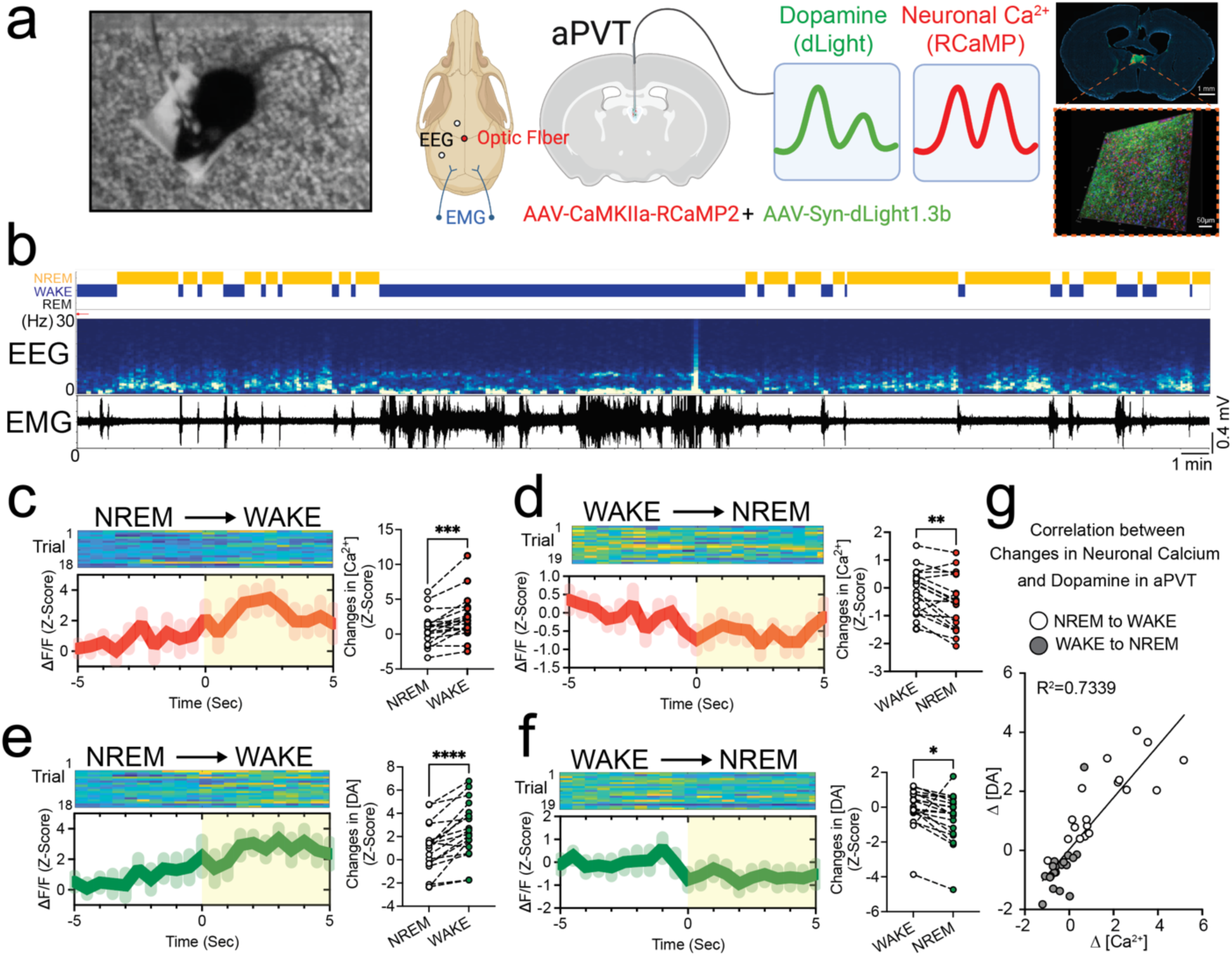
The neuronal calcium transients in the PVT are positively correlated with the transition of sleep pattern to Wake. (a) The placement of electrodes for EEG-EMG data collection and an optic-fiber for fiber-photometry. (b) Example traces of EEG-EMG recording. (c-g) Sorted data showing the simultaneous neuronal calcium (c-d) and dopamine (e-f) transients in the anterior part of paraventricular nucleus of thalamus (aPVT) as sleep patterns shift. (Figure 2c, Paired t-test, t=4.142, df=17, P= 0.0007, N=18/group), (Figure 2d, Paired t-test, t=3.766, df=18, 0.0014, N=19/group), (Figure 2e, Paired *t*-test, t=5.572, df=17, P <0.0001, N=18/group), (Figure 2f, Paired t-test, t=2.620, df=18, P=0.0174, N=19/group). *p<0.05, **p<0.01, ***p<0.001, ****p<0.0001. (e) Correlation between the changes in dopamine and neuronal calcium transients in the aPVT. Simple linear regression of Correlation. (Figure 2e, R^2^ = 0.7339, F = 96.54, DFn, Dfd = 1, 35, Y = 0.8779*X + 0.03826, P<0.0001).

### Withdrawal from repeated intermittent ethanol exposure increases neuronal activities in the PVT

To better understand the effects of withdrawal from repeated ethanol exposures on the PVT neurons, we used immunofluorescence staining with cFos, an immediate early gene marker of neuronal activity (Figure 3a) (Li et al., 2021; Zhang et al., 2022; Kooiker et al., 2023). We found that the CIE mice have significantly higher cFos expression than those in the ethanol naïve CON mice in the PVT, specially, the anterior part of the PVT (aPVT) (Figure 3b, Two-way ANOVA, F (1, 28) = 9.373, P=0.0048; CON vs CIE, aPVT p=0.0075, pPVT p=0.5055). This data suggests that the enhanced activity of the PVT neurons is accompanied with CIE-induced sleep disturbance and increased wakefulness in a sub-region dependent manner.

**Figure 3.**
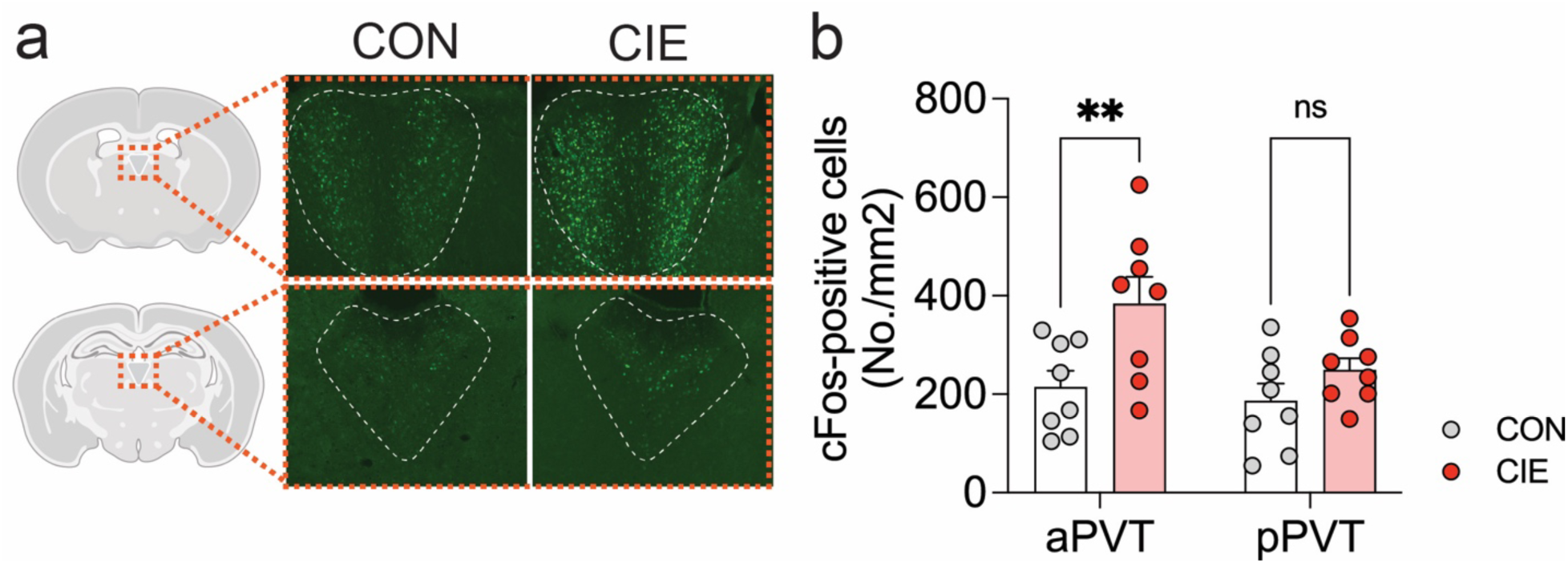
Withdrawal from repeated intermittent ethanol exposure increases neuronal activities in the PVT. (a-b) Representative figures (a) and pooled data (b) showing that cells expressing cFos, an immediate early gene, in the anterior PVT (aPVT) is increased after ethanol withdrawal. [Figure 3b: Two-way ANOVA, Interaction F (1, 28) = 1.988, P=0.1695, Region F (1, 28) = 4.610, P=0.0406, Group F (1, 28) = 9.373, P=0.0048, Bonferroni’s multiple comparisons test: CON vs CIE, aPVT: p=0.0075, pPVT p=0.5055]. N=8/group. Data represented as mean ± SEM. **p<0.01.

### Chemogenetic inhibition of aPVT neurons alleviates CIE-induced sleep disturbance

We observed that, in parrel to CIE-induced sleep disturbance including increased wakefulness, the aPVT neuronal activity is enhanced Figure 3). To further identify whether the PVT directly contributes to the sleep disturbance in ethanol-withdrawn mice, we sought to evaluate the CIE-induced sleep patterns when aPVT neuronal activity is inhibited (Figure 4a). For those, we infected the aPVT neurons with an adeno-associated viral vector serotype 5 (AAV5) expressing hM4Di, an inhibitory DREADD. This was under the control of a CaMKIIa promoter, which leads to selective expression in excitatory neurons (Figure b). Administration of JHU37160 (J60, 0.3 mg/kg, i.p.) significantly reduced the amount of cFos labeled cells in the aPVT of the hM4Di-injected CIE mice (Figure 4c, Unpaired t-test, t=2.841, df=6, P=0.0295). Interestingly, the chemogenetic inhibition of aPVT neurons increased the NREM and decreased the WAKE in the hM4Di-injected CIE mice compared to those in the vehicle-administered CIE mice (Figure 4d,e) [Two-way ANOVA, F (2, 18) = 14.96, P=0.0001, CIE+SAL vs CIE+60, WAKE: p=0.0038, NREM p=0.003, REM: p>0.999], indicating that the inhibition of PVT neurons could ameliorate the sleep disturbance induced by withdrawal from repeated ethanol exposure.

**Figure 4.**
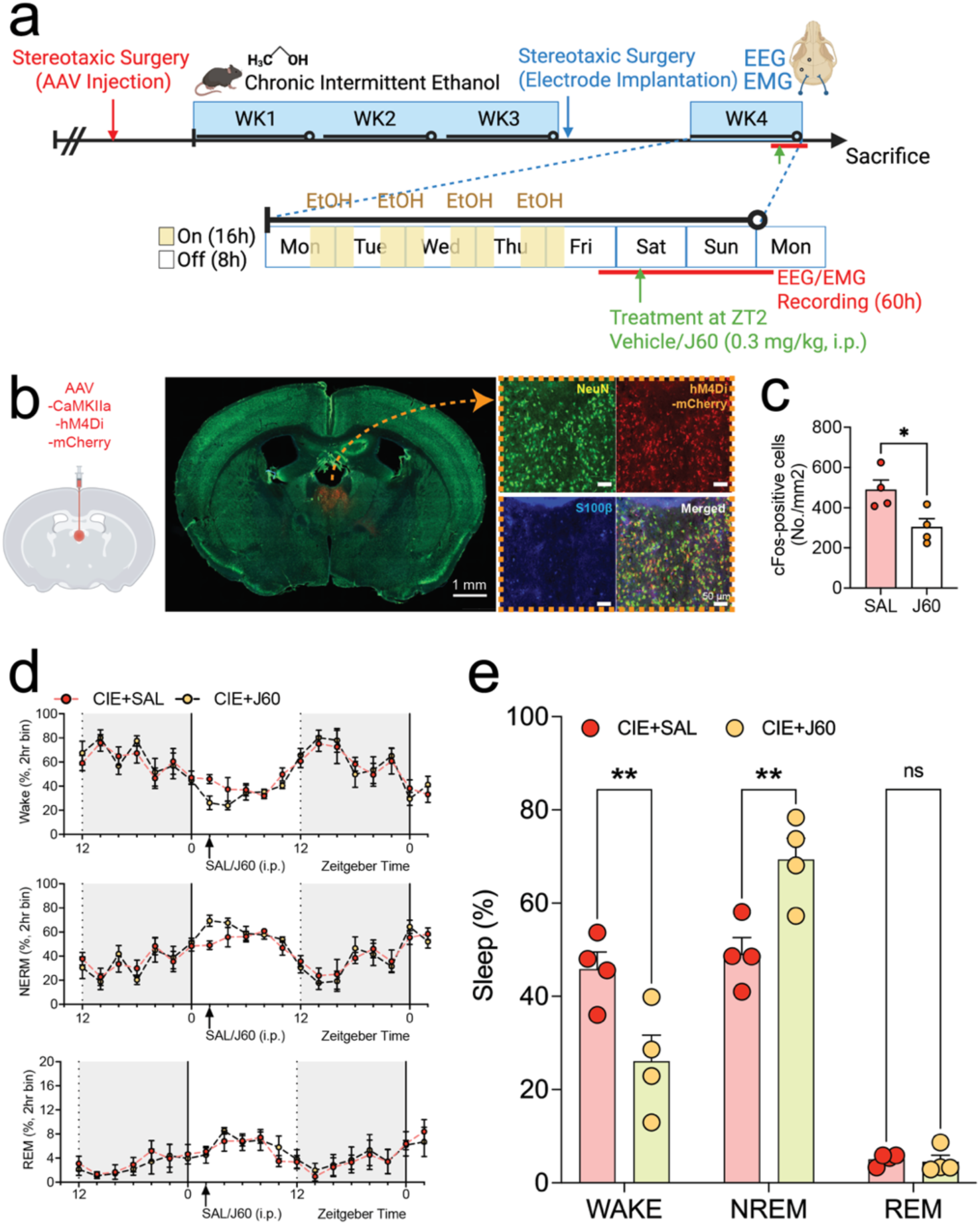
Chemogenetic inhibition of aPVT neurons alleviates the CIE-induced sleep disturbance. (a) Experimental schedule. (b) Representative figure showing the expression and function of hM4Di-mCherry in the aPVT. (c) Pooled data showing the reduced cFos-positive cells in the aPVT by J60 administration compared to those by vehicle (saline) administration. Unpaired t-test, t=2.841, df=6, N=4/group. *p<0.05. (d-e) Pooled data showing that the inhibition of aPVT neuronal activities by chemogenetic application rescues the CIE-induced sleep disturbance. N=4/group. Two-way ANOVA, Interaction F (2, 18) = 14.96, P=0.0001, Sleep type F (2, 18) = 111.2, P<0.0001, Group F (1, 18) = 7.771e-016, P>0.9999, Bonferroni’s multiple comparisons test: CIE+SAL vs CIE+J60, WAKE p=0.0038, NREM p=0.003, REM p>0.9999]. Data represented as mean ± SEM. *p<0.05, **p<0.01. J60: JHU37160 (0.3 mg/kg, i.p.).

### PVT show distinguishable characters in the expression and function of m-currents and the expression of KCNQ2 is reduced in the aPVT of CIE mice

The biochemical and chemogenetic results demonstrate that the PVT neuronal activity was increased in a subregion dependent manner specially alongside with the antero-posterior axis and the inhibition of the aPVT alleviated the wakefulness in Ethanol withdrawn mice. Previous work has shown there are genetically-, anatomically-, and functionally-distinct cell types in the PVT according to the genetic markers and/or locations along with medio-lateral or antero-posterior axis (aPVT vs. pPVT) (Gao et al., 2020; Shima et al., 2023). In the neuron, the afterhyperpolarization is mediated by three different ion channel subtypes: small-conductance calcium-activated K+ (SK) channels, voltage-dependent M channels (KV7/KCNQ), and the hyperpolarization-activated cyclic nucleotide-gated (HCN) channel (Gu et al., 2005; Dwivedi and Bhalla, 2021). Among them, M channels are one of the major channels that was not only affected by ethanol exposure (Kang et al., 2017; Cannady et al., 2018; Kang et al., 2019; Dwivedi and Bhalla, 2021), but also showed an important role to modulate sleep patterns (Li et al., 2022b; Whalley, 2022). Notably, we found that the protein expression of the M-channel subunit, KCNQ2 is significantly highly observed in the neurons of aPVT compared to those in the pPVT (Figure 5a,b; Unpaired t-test, t=4.751, df=8, P=0.0014), which is matched to the M-currents in the aPVT and pPVT (Figure 5c, Unpaired t-test, t=2.920, df=12, P=0.0128). To determine the effects of ethanol exposure on KCNQ2 expression in the aPVT, we performed western blotting to compare the ethanol naïve CON and CIE exposed groups (Figure 6a). We showed that KCNQ2 expression in the aPVT is significantly reduced in the CIE group compared to controls (Figure 6b, Unpaired t-test, t=2.920, df=12, P=0.0128).

**Figure 5.**
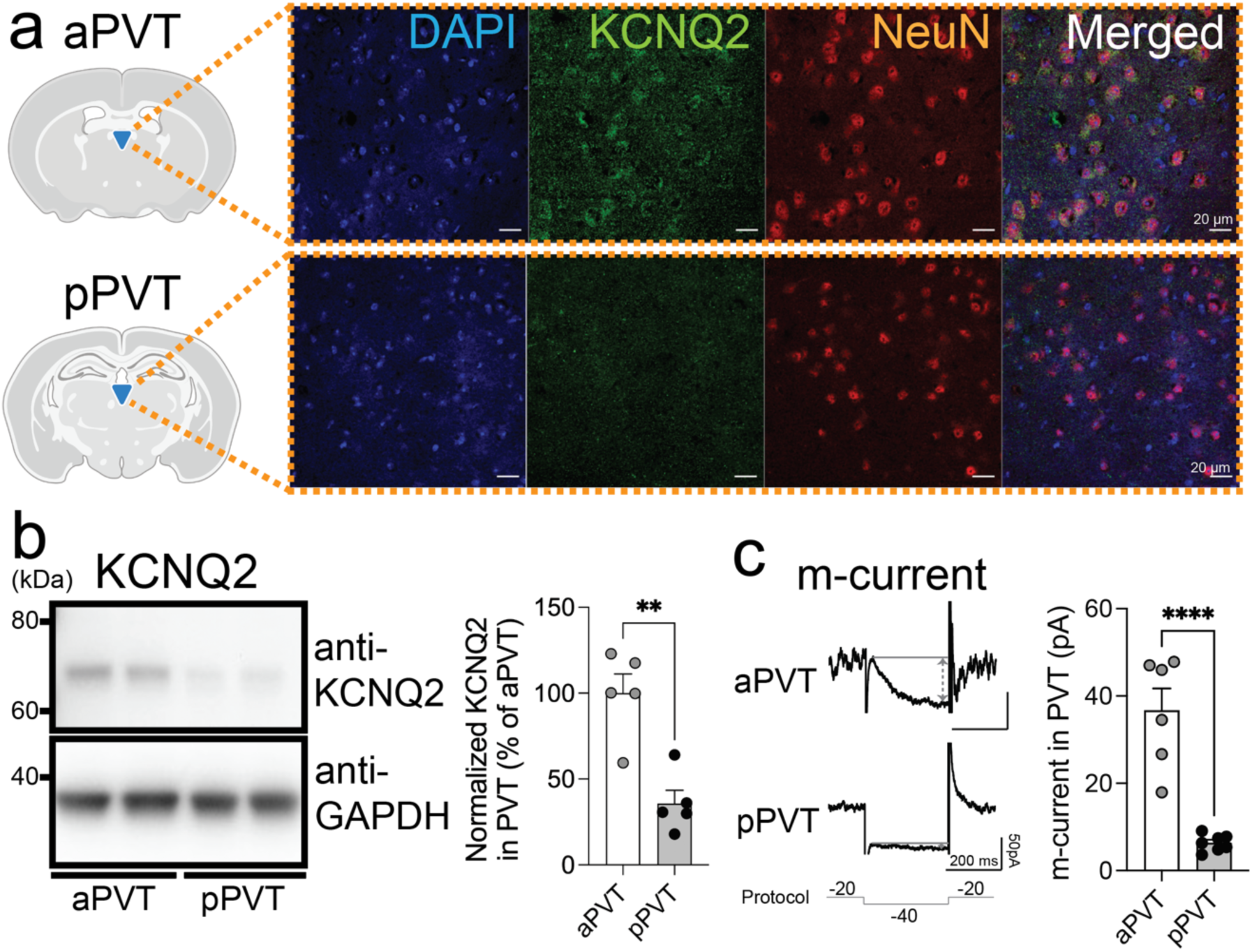
Anterior and posterior PVT show distinguishable characters in the expression and function of m-currents. (a) Representative immuno-labeling figures showing the expression of KCNQ2, a subunit of m-type potassium channel, and NeuN, a neuronal marker, in the anterior and posterior PVT. (b-c) Representative western blots of the expression of KCNQ2 (top) and GAPDH (bottom) and its pooled data (b) showing that KCNQ2 expression is much higher in the aPVT compared to that in the pPVT. Unpaired t-test, t=4.751, df=8, P=0.0014, N=5/group. Representative electrophysiological traces and pooled data (c) showing that the m-current is much higher in the aPVT compared to that in pPVT. Unpaired t-test, t=6.407, df=11, P<0.0001, N=6-7/group. Data represented as mean ± SEM. **p<0.01, ****p<0.0001.

**Figure 6.**
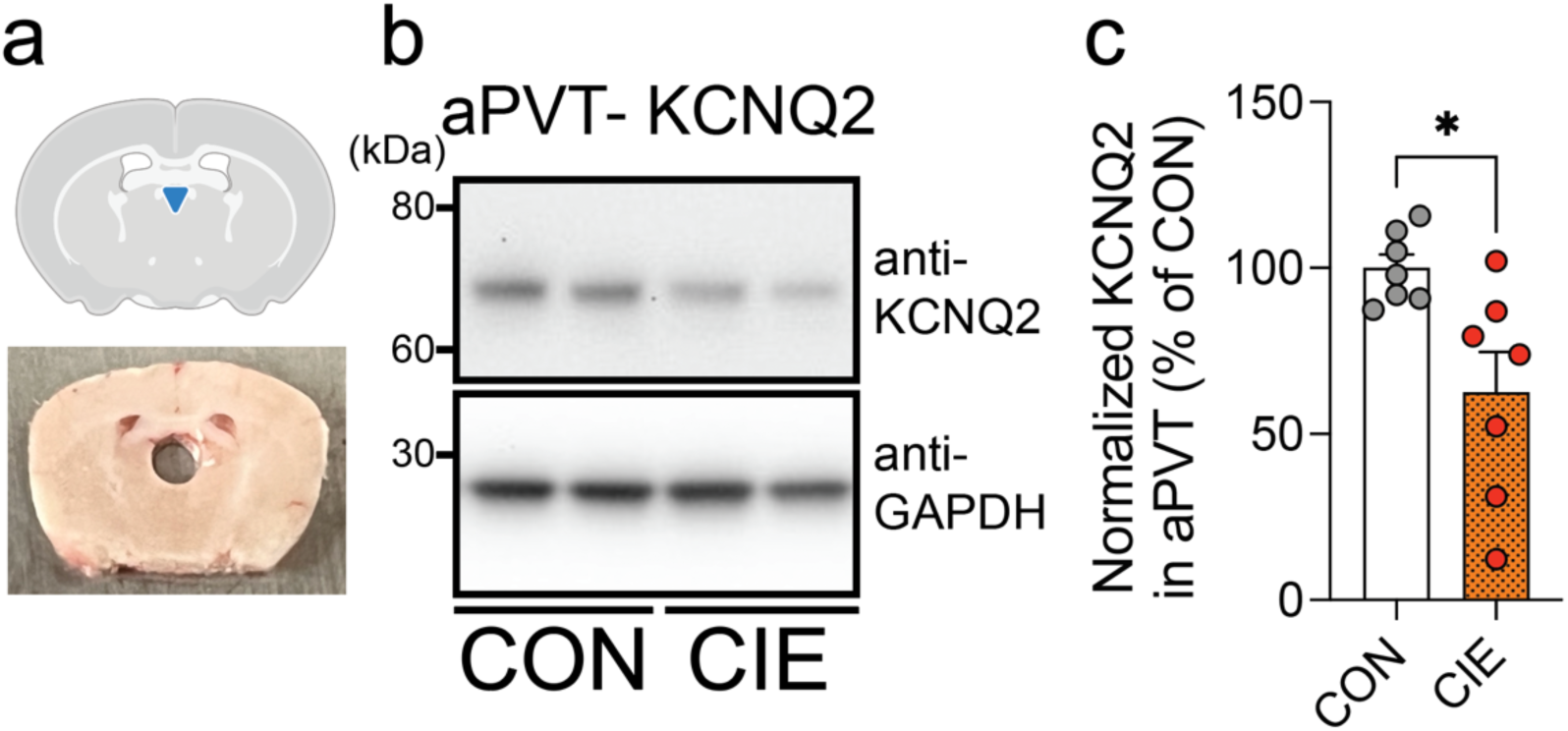
The expression of KCNQ2 is reduced in the aPVT of CIE mice. (a-c) Diagram and example of punched tissue (a), representative expression of KCNQ2 (right top, b) and GAPDH (right bottom, b) and, pooled data (c) of western blots showing that the KCNQ2 expression in the aPVT of CIE mice is significantly reduced compared to that of the CON mice. N=7/group. Unpaired *t*-test. Data represented as mean ± SEM. *p<0.05.

### Rescue of KCNQ2 expression in the aPVT ameliorates the CIE-induced sleep disturbance

To directly test whether rescue of M-channel in the PVT alleviates the sleep disturbance in CIE mice, we measured the sleep patterns after overexpression of KCNQ2 in the aPVT (Figure 7a). We rescued the KCNQ2 expression by co-injecting an AAV expressing Cre recombinase under hSyn promotor with an AAV expressing KCNQ2 Cre-dependently under an Ef1a promotor into the aPVT (Figure 7b), and found that the NREM was significantly increased, and the WAKE time was significantly decreased in the CIE mice with the KCNQ2 overexpression in the aPVT compared to those of CIE mice that just received the same stereotaxic procedures (Figure 7c-d) [Z0-12: Two-way ANOVA, F (2, 54) = 11.30, P<0.0001, CIE+CON vs CIE+Q2, WAKE: p=0.0055, NREM p=0.0034, REM: p>0.999], (Figure S2) [Z12-24: Two-way ANOVA, F (2, 54) = 0.7145, P<0.4940, CIE+CON vs CIE+Q2, WAKE: p>0.9999, NREM p>0.9999, REM: p>0.9999]. These data suggest that the restoration of KCNQ2 expression in the aPVT mitigates the sleep disturbances induced by CIE exposure.

**Figure 7.**
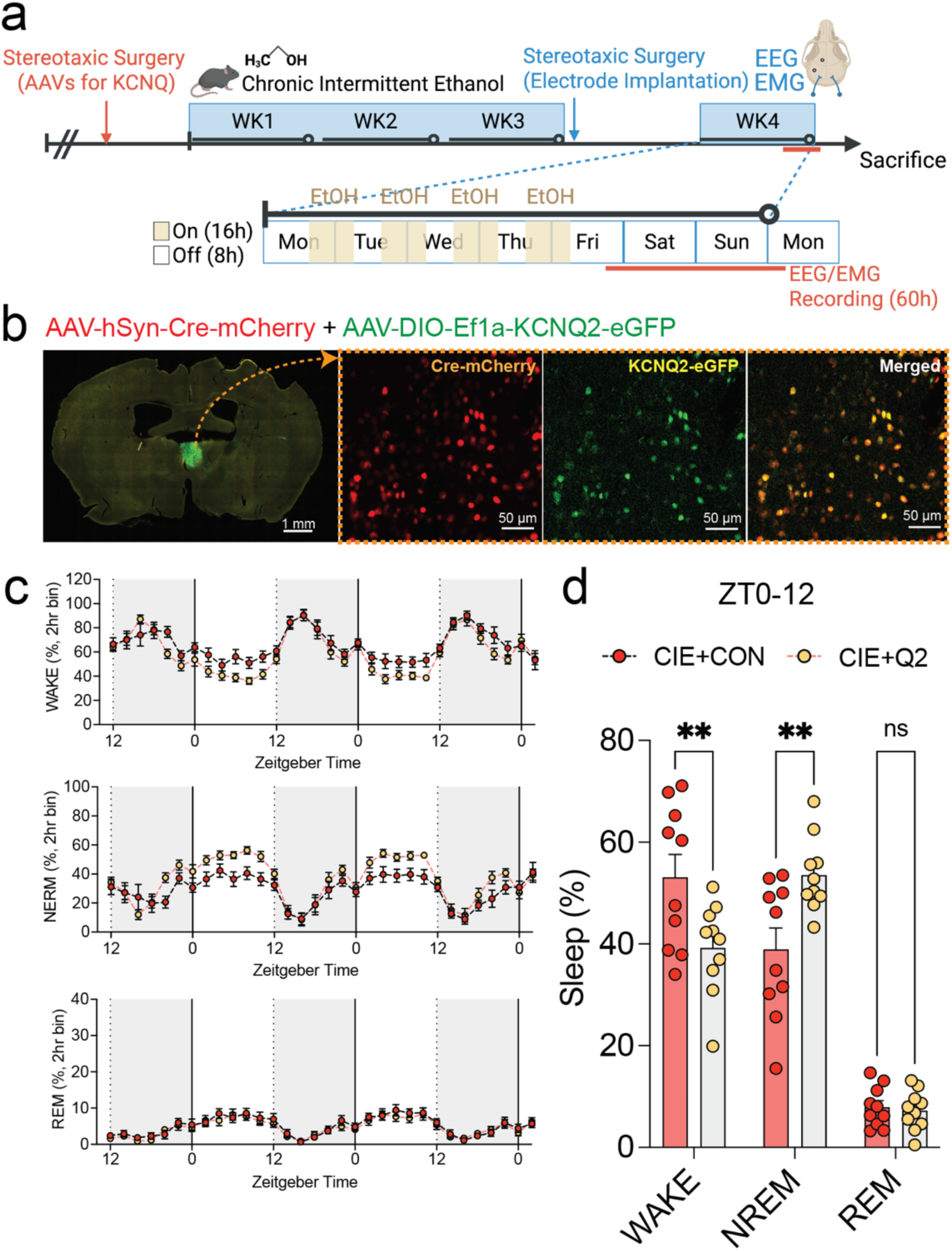
Rescue of KCNQ2 expression in the aPVT ameliorates the CIE-induced sleep disturbance. (a) Experimental schedule. (b) Representative figure showing the expression of KCNQ2 in the neurons of aPVT. (c-d) Pooled data showing that KCNQ2 overexpression in the aPVT of CIE mice ameliorates the CIE-induced sleep patterns in the day light cycle. [Figure 7d, Two-way ANOVA, Interaction F (2, 54) = 11.30, P<0.0001, Sleep type F (2, 54) = 111.0, P<0.0001, Group F (1, 54) = 1.678e-014, P>0.9999, Bonferroni’s multiple comparisons test: CIE+CON vs CIE+Q2, WAKE p=0.0055, NREM p=0.0034, REM p>0.9999]. N=10/group. Data represented as mean ± SEM. **p<0.01.

Taken together, our findings support the hypothesis that withdrawal from chronic ethanol exposure enhanced the PVT neuronal activities in an antero-posterior axis dependent manner, leading to the enhanced wakefulness during the withdrawal, at least partly, via the downregulation of M-channels in the anterior PVT.

## Discussion

In this study, we demonstrated that the neuronal adaptation in the PVT altered sleep patterns in mice withdrawn from chronic intermittent ethanol exposure. Specifically, silencing anterior PVT (aPVT) excitatory neurons attenuated the increased Wakefulness in CIE mice. We also confirmed our recent finding that M-channel subunit, KCNQ2, in the aPVT was reduced during CIE. Furthermore, we showed that genetic upregulation of M-channels in the aPVT alleviated the sleep disturbance including the increase in Wakefulness in CIE mice. The findings indicate that the downregulation of M-channels in the aPVT of CIE mice leads to the hyperactivity of aPVT neurons and disrupts sleep. Consequently, the PVT is crucial to the processing of brain adaptations associated with sleep and repeated alcohol.

We also identified molecular adaptations in the PVT neurons of CIE mice. Ion channels are important modulators of neuronal excitability (Catterall, 1984; Greene and Hoshi, 2017). Among the components, potassium ion channels act as a break to prevent excessive firing (Cooper and Jan, 2003; Greene and Hoshi, 2017). Consistent to the previous studies showing the reduced expression of M-channel subunits, KCNQ family, in diverse brain regions after the chronic ethanol exposed rodents compared to those in naïve counterparts and behavioral rescue by the enhancement of M-currents (Knapp et al., 2014; Kang et al., 2017; Cannady et al., 2018; Kang et al., 2019), this study demonstrated that KCNQ2, which are highly expressed in the aPVT relative to the pPVT, were diminished in CIE mice, along to sleep disturbances and this abnormal sleep patterns observed in CIE mice was attenuated by AAV-driven overexpression of KCNQ2 in the aPVT. Although M-channels have been identified as a modulator of neuronal excitability and sleep patterns (Cooper et al., 2001; Li et al., 2022b), our study is, to the best of our knowledge, the first study confirming the involvement of M-channels in alcohol-induced sleep disturbances.

Recently updated human brain imaging and animal studies have shown that the PVT undergoes structural and functional changes in parallel with sleep cycles and long-lasting sleep disturbance (Colavito et al., 2015; Ren et al., 2018; Ahrens and Ahmed, 2020). Sleep disturbance is thought to be caused by aberrant neuronal responses in the process of homeostatic sleep drive and circadian alerting signals (Novak et al., 2000; Nascimento et al., 2008; Phan and Malkani, 2019; Tossell et al., 2023; Ma et al., 2024). A major contribution of this study is the identification of PVT excitatory neurons as regulators of sleep modulation specially affected by chronic ethanol exposure. Our observation that silencing PVT excitatory neurons reduced Wakefulness is consistent with recent rodent study to evaluate diverse external-stimuli induced changes in sleep patterns (Ren et al., 2018).

Notably, morphological, behavioral and electrophysiological evidence suggests that several neurotransmitters are involved in the modulation of arousal and sleep cycle under both physiological and pathological conditions (Sardi et al., 2018; Antila et al., 2022; Toth and Burgess, 2025). In the brain, dopaminergic neurons in hypothalamic area and ventral tegmental area (VTA) have been considered an important component of modulation of sleep cycles triggered by external stimuli, or internal circadian rhythm, or interest/salience (Dzirasa et al., 2006; Liu et al., 2017; Toth and Burgess, 2025). The PVT not only has a dense expression of a series of dopaminergic receptors in a subregion-specific manner (Gao et al., 2020; Penzo and Gao, 2021; Shima et al., 2023), but receives multiple projections from diverse brain regions plausibly releasing dopamine (Beas et al., 2018; Otis et al., 2019; Guo et al., 2023). Given the role of dopamine signaling as a fine tuner of sleep patterns in brain (Hasegawa et al., 2022) and its modulation of PVT related behaviors (Beas et al., 2018), Our findings that dopamine transients correspond with PVT neuronal activity and subsequent alterations in sleep patterns, implies the importance of dopaminergic signals in the PVT for behavioral fine-tuning in sleep cycles. Thus, it will be intriguing to evaluate how dopamine signals influence the PVT activities and consequent behaviors according to activation of molecularly distinct dopaminergic receptor-positive PVT neurons. Although several major studies show the activation of PVT neurons induce wakefulness (Ren et al., 2018; Li et al., 2022a), there are also disparate reports showing the reduced wakefulness according to the activation of subpopulations of PVT neurons (Matyas et al., 2018; McGinty and Otis, 2020). This discrepancy may arise from the cellular molecular and structural diversity in the PVT, suggesting unmet needs to study the molecularly distinct cell types. Understanding how those characteristics are altered by withdrawal from repeated ethanol exposure is crucial for comprehending the detailed brain activity associated with alcohol withdrawal-induced sleep disturbances. Furthermore, considering the signal cascade that dopamine receptor type 1 (D1R) can block M-channels through the Erk signaling pathway (Tsuboi et al., 2022), these dopamine transients could be an important factor to modulate the activity of M-currents within the circuit, hence affecting sleep regulation. Regarding the different expression of dopaminergic receptors in the PVT, it would be intriguing to explore how dopamine signals affect the PVT subpopulations via the activity of M-channels. Further studies are needed to test these possibilities.

Given that M-channels are slowly closing at close-to-resting membrane potentials (RMP) and modulating afterhyperpolarization (AHP) (Gu et al., 2005; Hu et al., 2007; Tzingounis and Nicoll, 2008; Brown and Passmore, 2009; Varghese et al., 2023) and that repeated alcohol exposure reduce the M-channels in brain (McGuier et al., 2015; Kang et al., 2017; Kang et al., 2019), M-channels may play an important role mainly in the aPVT excitatory neuronal adaptations during alcohol withdrawal. However, there is a possibility that other potassium channels or the activities of KCNQ family in non-neuronal cells are also involved in the PVT pathological adaptation. A recent study has demonstrated several ion channels that have a different expression in the PVT in a cell-type dependent manner alongside the anteroposterior axis (Shima et al., 2023). For example, Kir4.1, one of the inwardly rectifying potassium channels that is predominantly localized in non-neuronal glial cells, astrocytes and reduced by withdrawal from repeated ethanol exposure, which is similar paradigm to this study (Ayers-Ringler et al., 2016), has been known to be modulating extracellular potassium levels and indirectly affecting neighbor neuronal firing by the modulation of potassium buffering (Nwaobi et al., 2016; Cui et al., 2018; Kinboshi et al., 2020). In addition, current studies looking into the role of the KCNQ family, particularly KCNQ2-5, in neurological disorders have focused on their function in neurons. Recent research has revealed that KCNQ channels play a significant role in astrocytes as a key regulator of neighbor neuronal excitability via controlling neurotransmitters (Graziano et al., 2024; Xia et al., 2024). Therefore, it will be of interest to further investigate the changes in functions of other potassium channels, including other subunits of KCNQ and its co-factors localized in the PVT during alcohol withdrawal.

It is still unclear whether alcohol exposure modifies sleep patterns through the PVT or whether changes in sleep patterns lead to additional alcohol-induced brain adaptation. Several studies report that, when mice are exposed to repeated intermittent ethanol exposure, mice underwent brain adaptation within one to two weeks. However, according to our data, the mice didn’t show the sleep disturbance withdrawal from repeated ethanol exposure for one week, suggesting that repeated alcohol exposure and withdrawal seem to alter sleep patterns. To understand this better, in future studies, additional experiments will involve activating the PVT to produce alterations in sleep patterns and subjecting the mice to the alcohol voluntary drinking paradigm to determine whether sleep disturbances caused by PVT hyperactivity result in increased alcohol consumption.

We modulated the activity of PVT neurons in freely moving mice by AAV-mediated gene transfer for the expression of hM4Di DREADDs in PVT neurons, coupled with systemic injection of the ligand JHU37160 (J60, 0.3 mg/kg, i.p.). Notably, although this new generation of DREADDs agonist, J60, has a very high in vivo DREADD potency in CNS, diverse responses such sedative effect including reduced locomotion (10 mg/kg, mice) and anxiogenic effect (1 mg/kg, rat) were observed when administered as a high dose (Van Savage and Avegno, 2023; Aomine et al., 2024). Further control validations such as dose-dependent responses and more control groups such as low-dose clozapine treatment should be performed in future studies.

In this study, we have provided several lines of evidence regarding the impact of the PVT on sleep disturbance in mice withdrawn from chronic intermittent ethanol exposure. Wakefulness is heightened in ethanol-withdrawn animals during withdrawal from chronic ethanol exposure, in parallel with elevated basal activity of aPVT neurons and down-regulation of M-channels. Chemogenetic inhibition or overexpression of the M-channel subunit KCNQ2 in aPVT neurons mitigates sleep disturbances, including heightened wakefulness. The data collectively indicate that M-channels significantly contribute to the hyperactivity of PVT neurons in ethanol-withdrawn mice, and PVT activity is essential for sleep disturbances. Modulating region-specific PVT activity, particularly via M-channels, could be therapeutically beneficial for addressing sleep disturbances and alcohol use disorder.

## Data availability

All data are available from the authors upon reasonable request.

## Acknowledgment

We especially thank to Drs. Karl Deisseroth, Lin Tian, and Bryan Roth for providing plasmid DNA. We thank Dr. Mungyu Song, Kishan Trivedi, Sahil Patel, Lesley Barksdale, Ni Na Ngo, Kaleb Louisy, Bansari Patel, and Chloe Yuri Woo for technical assistance and critical discussion. We express appreciation to all laboratory members for their valuable discussions and comments. Some of the figures were created with BioRender.com. This research was supported by the National Institute of Health (AA027773, MH137204 to SK).

## Author contributions

SK and AB designed the study. SK, AB, and HK performed all behavioral, electrophysiological, and biochemical experiments. SK and AB performed the stereotaxic surgeries. SK, AB, JL, VK, DJ, and HK collected, processed, and imaged tissue for histology. SK, AB, and JL performed the collection and analysis of the EEG/EMG data. AB, VK, HK, and SK wrote the manuscript. All authors reviewed and edited the manuscript.

---

**Figure S1.**
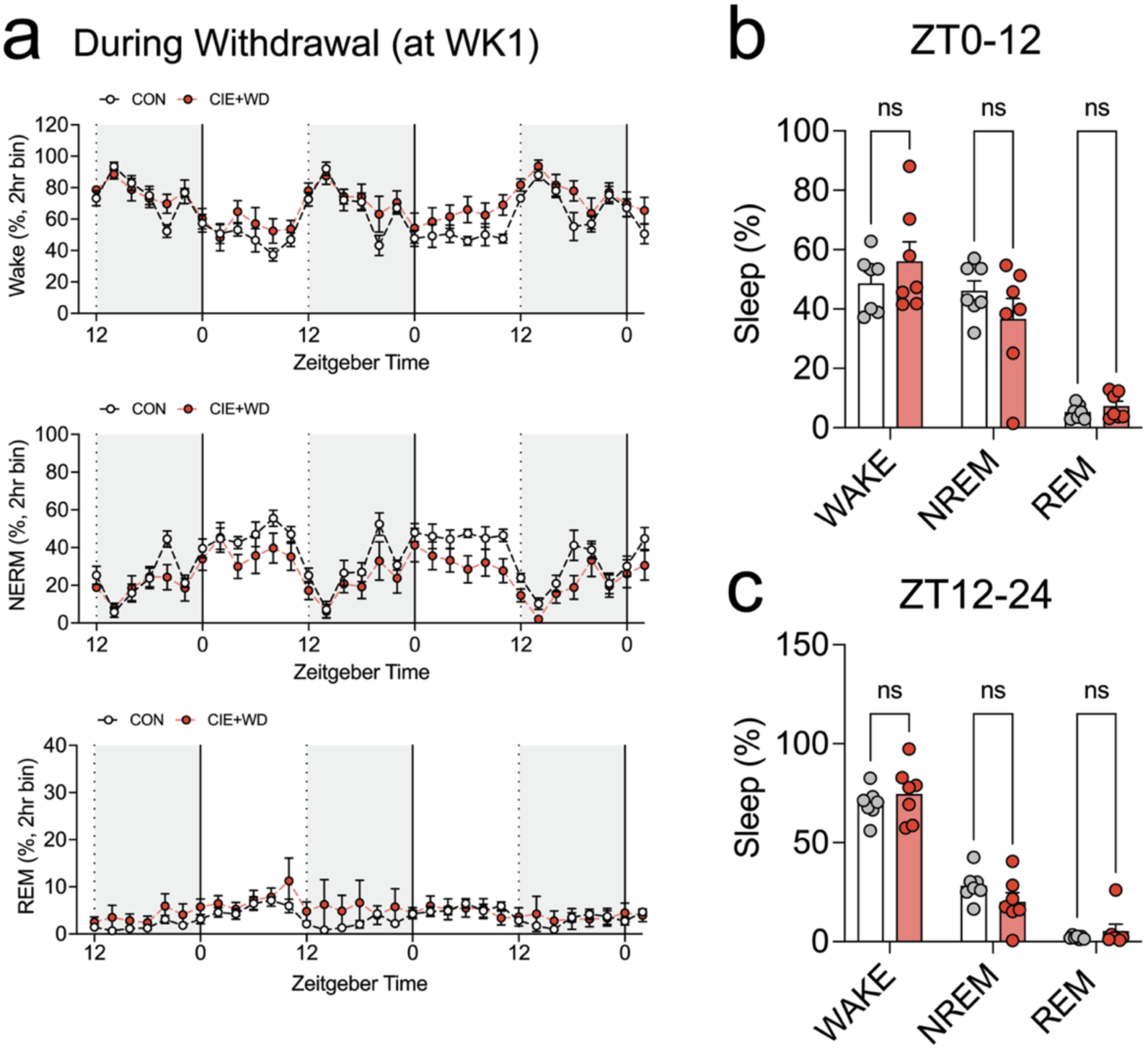
Evaluation of the changes in sleep patterns at 1 day withdrawal from repeated ethanol exposures for 1 week. (a-c) Representative figures (a) and pooled data (b-c) showing that no significant difference is observed during withdrawal from chronic intermittent ethanol exposure for 1 week. [Figure S1b, Interaction F (2, 36) = 1.844, P=0.1728, Sleep type F (2, 36) = 57.80, P<0.0001, Group F (1, 36) = 7.914e-016, P>0.9999, Bonferroni’s multiple comparisons test: CON vs CIE-WD, WAKE p=0.7513, NREM p=0.4349, REM p>0.9999], [Figure S1c, Interaction F (2, 36) = 1.903, P=0.1639, Sleep type F (2, 36) = 185.0, P<0.0001, Group F (1, 36) = 1.370e-014, P>0.9999, Bonferroni’s multiple comparisons test: CON vs CIE+WD, WAKE p>0.9999, NREM p=0.3676, REM p>0.9999]. N=7/group. Data represented as mean ± SEM.

**Figure S2.**
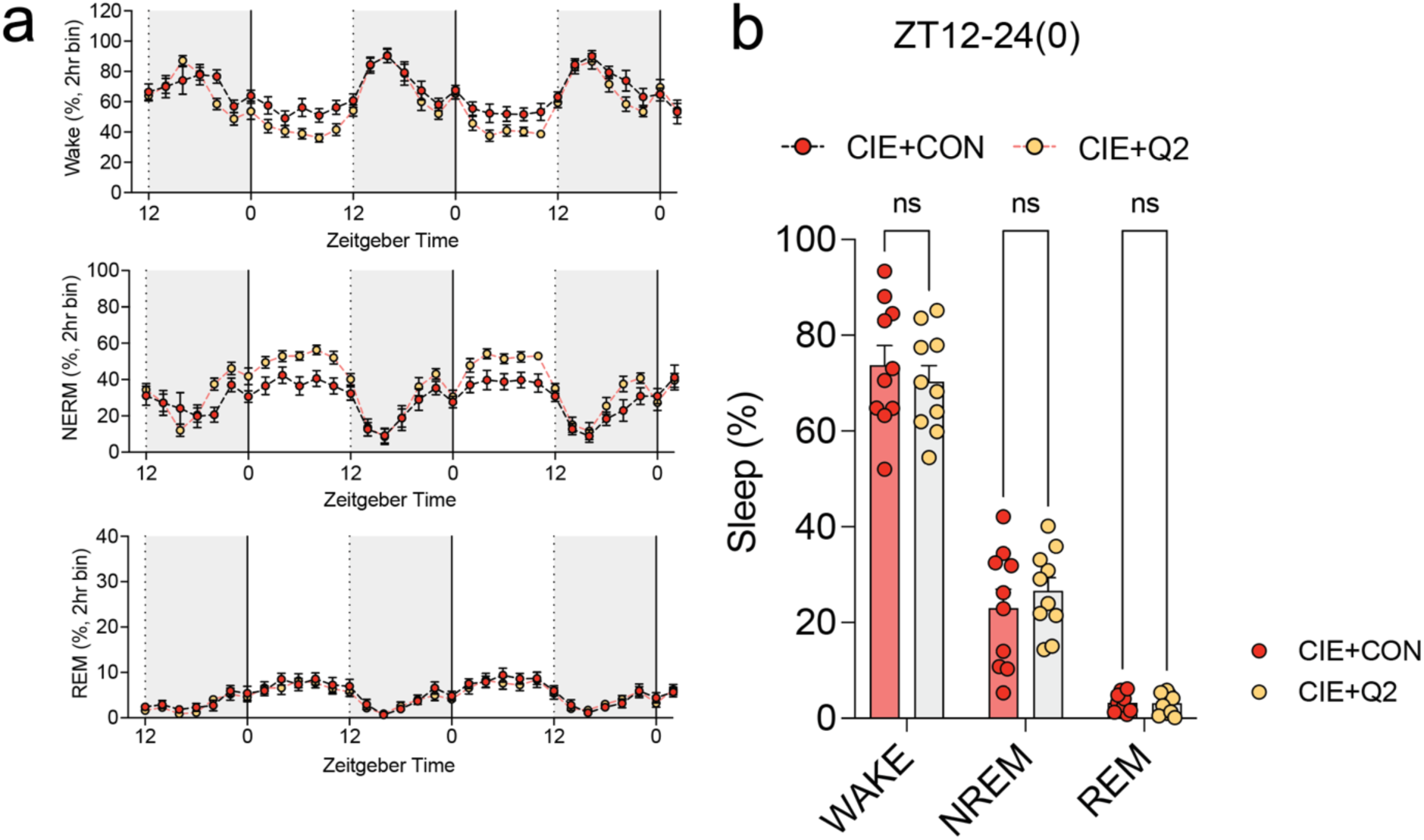
Evaluation of the rescue of KCNQ2 expression in the aPVT in the dark light cycle. (a-b) Pooled data showing that no significant difference is observed in the dark light cycle after the overexpression of KCNQ2 in the aPVT of CIE mice. [Figure S2d, Two-way ANOVA, Interaction F (2, 54) = 0.7145, P=0.4940, Sleep type F (2, 54) = 287.1, P<0.0001, Group F (1, 54) = 1.495e-014, P>0.9999, Bonferroni’s multiple comparisons test: CIE+CON vs CIE+Q2, WAKE p>0.9999, NREM p>0.9999, REM p>0.9999]. N=10/group. Data represented as mean ± SEM.

